# Developmentally programmed changes in cytoplasmic mechanics revealed by active microrheology in *C. elegans* embryos

**DOI:** 10.64898/2026.05.19.726147

**Authors:** Saaya Koizumi, Ayama Tokuyasu, Akinori W. M. Miyamoto, Takayuki Torisawa, Hirokazu Tanimoto, Akatsuki Kimura

## Abstract

Cytoplasmic mechanical properties are often treated as constant background parameters, yet whether they change systematically during development remains unclear. Here, we directly measured cytoplasmic mechanics during early embryogenesis of *Caenorhabditis elegans* by establishing active microrheology using micrometer-sized magnetic droplets. Active microrheology revealed a progressive decrease in creep compliance from the 1-cell to the 8-cell stage, indicating a progressive stiffening of the local cytoplasmic environment during development. This decrease persisted even when cytokinesis was inhibited, demonstrating that it cannot be explained solely by geometric changes associated with cell division. Passive microrheology using 40-nm fluorescent beads showed a consistent decrease in probe mobility over development. Together, these results demonstrate that cytoplasmic mechanical properties undergo a gradual, developmentally programmed change during embryogenesis that cannot be explained by cell division–associated geometry alone.

## Introduction

The cytoplasm is a crowded, heterogeneous, and energy-consuming medium whose physical properties are central to cellular function. Its viscosity and elasticity influence processes ranging from molecular diffusion and reaction kinetics to organelle transport, positioning, and mechanical stability (Luby-Phelps, 2000; Parry et al., 2014; Delarue et al., 2018; Garzon-Coral et al., 2016; Najafi et al., 2023). At the molecular scale, cytoplasmic crowding modulates diffusion and reaction rates, whereas at the micron scale, viscous drag and elastic restoring forces shape the dynamics and positioning of structures such as nuclei and spindles (Persson et al., 2020; Duch et al., 2020; Xie et al., 2022).

In many experimental and theoretical frameworks, cytoplasmic mechanical properties are treated as time-independent background parameters that provide a constant physical environment for intracellular processes (Garzon-Coral et al., 2016; Anjur-Dietrich et al., 2024; Xie et al., 2025; Sun et al., 2025). This view is consistent with the expectation that cytoplasmic viscosity remains relatively stable to support robust cellular function. However, accumulating evidence indicates that cytoplasmic mechanical properties can undergo transient changes in specific biological contexts. For example, intracellular mechanics have been reported to vary during the cell cycle and differentiation (Taubenberger et al., 2020; Kickuth et al., 2026; McAndrews et al., 2014; Chen et al., 2016). In early embryogenesis, a temporally restricted increase in cytoplasmic viscosity has been observed during late 2-cell interphase in mouse embryos, suggesting stage-specific modulation (Ye and Homer, 2024). Similarly, dramatic but reversible increases in cytoplasmic viscosity have been reported in specialized cellular states such as yeast spores (Sakai et al., 2024). Together, these observations suggest that while cytoplasmic viscosity is often treated as constant, it can exhibit transient and context-dependent modulation.

During early embryogenesis, cells undergo rapid and successive divisions without growth, resulting in a stepwise reduction in cell size and systematic changes in geometric constraints such as volume and boundary conditions. Cytoplasmic mechanical properties have been shown to change in response to experimental manipulation of cell size (Tan et al., 2025). Whether and how similar regulation occurs during physiological cell size reduction in embryogenesis remains unclear. Under physiological conditions, changes in the behavior of large intracellular structures have been observed during embryogenesis. For example, in sea urchin embryos, the stability of spindle positioning increases with successive cell divisions (Xie et al., 2025). This phenomenon has been attributed to increased friction between the spindle and the cell boundary, rather than changes in cytoplasmic viscosity (Xie et al., 2021; Xie et al., 2025). Similarly, magnetic-tweezer measurements in *C. elegans* embryos have shown that spindle mobility is reduced at the 2-cell stage compared to the 1-cell stage, primarily due to interactions between the spindle and the cell cortex, without major changes in cytoplasmic viscosity (Garzon-Coral et al., 2016). However, direct experimental examinations of the material properties of the cytoplasm itself across developmental stages during embryogenesis have remained scarce. As a result, it is still unclear whether the material properties of the cytoplasm change during development, and if so, to what extent such changes contribute to the mechanical behaviors of intracellular structures.

Addressing this question requires methods that can directly measure cytoplasmic mechanical properties under well-defined conditions. Previous studies have often relied on passive microrheology (PMR), which infers mechanical properties from spontaneous fluctuations (Tseng et al., 2002; Weihs et al., 2007; Crocker et al., 2007). However, PMR cannot distinguish changes in driving forces from changes in material resistance, and is particularly limited in the cytoplasm where fluctuations are dominated by active, nonequilibrium processes (Ebata et al., 2023). In addition, cytoplasmic mechanical properties depend on the observation length scale, and measurements at the micrometer scale relevant to large intracellular structures remain scarce.

Active microrheology (AMR), which applies defined external forces and measures the resulting mechanical response, provides a complementary approach that overcomes these limitations (Ahmed et al., 2015; Vos et al., 2024; Fernandez et al., 2025). Among the available AMR approaches, magnetic tweezers uses magnetic probes and an external magnetic field to apply forces (Crick and Hughes 1950, Yagi 1961, Hiramoto 1969). Magnetic tweezers have been widely used to measure intracellular mechanical properties due to their high specificity and minimal perturbation to surrounding cellular structures (Wang et al., 2022; Kongari et al., 2025). To probe cytoplasmic mechanics under defined forces, sufficiently large probes are required to generate measurable displacements. Such micrometer-sized probes are also advantageous, as they approximate the scale of large intracellular organelles and therefore report on mechanically relevant length scales within the cell. However, implementing AMR in living cells has been technically challenging, as direct injection of large solid beads typically requires thick needles, which can cause substantial damage to the sample. The use of ferrofluids as deformable probes has enabled the formation of micrometer-sized droplets inside cells following injection through fine needles (He et al., 2014; Doubrovinski et al., 2017; Orii and Tanimoto, 2025), expanding the applicability of magnetic tweezer–based AMR in living systems.

Here, we investigate how cytoplasmic mechanical properties change during early embryogenesis in *C. elegans* using PMR and AMR. We ask whether cytoplasmic mechanical properties change systematically over developmental time, and whether they can be explained by geometric factors such as cell-size reduction or instead reflect intrinsic, developmentally programmed changes in the cytoplasmic state. By addressing these questions, this study establishes cytoplasmic material properties as a dynamically regulated layer of developmental control.

## Results

### Passive microrheology reveals a progressive decrease in cytoplasmic mobility during early embryogenesis

To investigate how the physical properties of the cytoplasm change during development, we analyzed the motion of probe particles in *C. elegans* embryos using passive microrheology (PMR). In 1-cell-stage embryos, previous PMR studies using 40 nm tracer beads (Torisawa et al., 2025) and 100 nm tracer beads (Daniels et al., 2006) reported diffusion coefficients in the range of approximately 0.01–0.1 μm^2^/s and MSD power-law scaling exponents close to 1, consistent with nearly diffusive motion on this scale. Here, we examined how these nanoscale mobility parameters change during early embryogenesis. 40-nm fluorescent beads were injected into the gonad and subsequently incorporated into embryos through normal oogenesis after a period of incubation. Because the beads were not directly injected into embryos, this approach enabled non-invasive incorporation into developing embryos. Embryos containing fluorescent beads were imaged by confocal microscopy. Time-lapse imaging were acquired separately at each developmental stage from the 1-cell to the 8-cell stage, with images acquired every 30 ms at a single focal plane. Individual beads were tracked from these time-lapse images.

For each developmental stage (1-, 2-, 4- and 8-cell stages; representative fluorescence images are shown in Fig. 1A), the trajectories of individual beads were used to calculate the mean squared displacement (MSD). The MSD decreased progressively from the 1-cell stage to the 8-cell stage (Fig. 1B), indicating that the cytoplasmic mobility becomes increasingly restricted during embryogenesis. The MSD exhibited a power-law dependence on the time interval (τ), described as MSD ∝ τ^α^. The scaling exponent α was approximately 1 across all developmental stages examined consistent with the behavior expected for purely viscous fluids (Fig. 1C). Given that α was close to 1, we fitted the MSD to a linear relation, MSD = 4Dτ, to estimate the diffusion coefficient D (Fig. 1C). While α remained nearly constant across stages, the diffusion coefficient D decreased progressively as embryogenesis proceeded (Fig. 1C), indicating that the reduction in MSD primarily reflects a decrease in effective diffusivity. Using the estimated diffusion coefficients, we calculated the apparent cytoplasmic viscosity based on the Stokes–Einstein relation (Fig. 1D). The viscosity was approximately two orders of magnitude higher than that of water, consistent with previous measurements using the same 40-nm beads (Torisawa et al., 2025), and lower than estimates obtained with larger probes (Daniels et al., 2006). Together, these results indicate that, within the probed timescale, cytoplasmic mechanics are dominated by viscous behavior and become progressively more resistive to diffusion during early embryogenesis.

**Figure 1.**
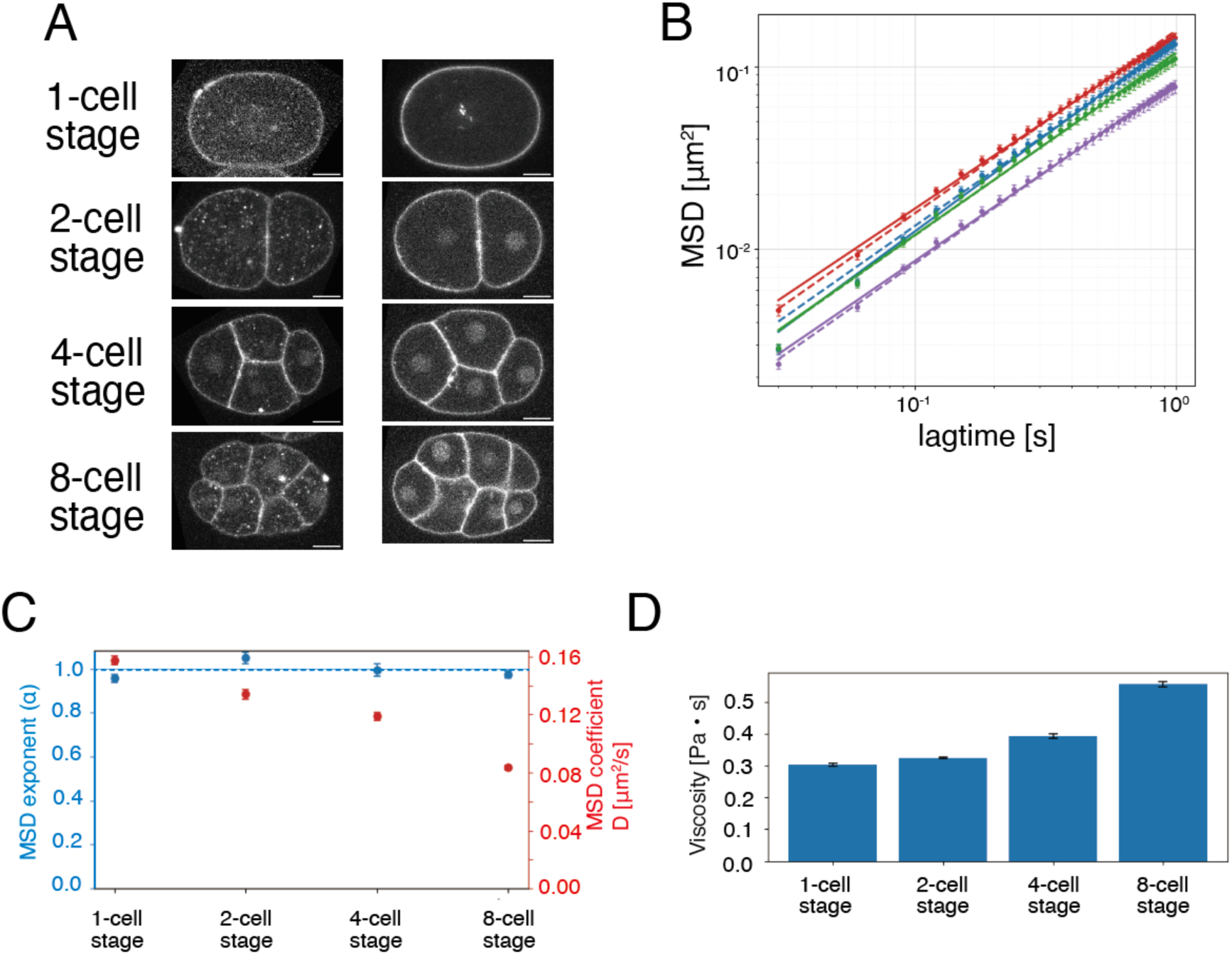
Passive microrheology shows that 40nm-diameter bead mobility decreases as *C. elegans* embryos develop from the 1-cell to the 4-cell stage. (A) Representative fluorescence images of 1-cell, 2-cell, and 4-cell stage *C. elegans* embryos of strain CAL2895 (mCherry::PH, mCherry::histone), shown with (left panels) and without (right panels) injection of 40 nm diameter fluorescent beads. (B) Mean square displacement (MSD) of fluorescent beads within embryos at each cell stage, calculated from bead movements. Color coding for embryo stages: 1-cell stage in red, 2-cell stage in blue, 4-cell stage in green, and 8-cell stage in purple. Fitted curves are overlaid using two models: a power-law fit, MSD = Cτ^α^ (dashed lines), and a linear fit, MSD = Dτ with α fixed at 1 (solid lines). Sample sizes were as follows: 1-cell stage, N = 5 embryos, n = 257 tracks; 2-cell stage, N = 5 embryos, n = 221 tracks; 4-cell stage, N = 5 embryos, n = 252 tracks; 8-cell stage, N = 6 embryos, n = 207 tracks. Error bars represent the standard error of the mean (SEM) of the MSD at each lag time, calculated as SEM = SD/√n, where SD is the standard deviation of squared displacements contributing to that lag time and n is the number of independent measurements. (C) Blue circles indicate the fitted scaling exponent α, and red circles indicate the fitted coefficient D in the relation MSD = Dτ for each cell stage, calculated from the MSD data in (B); error bars represent the 95% confidence intervals. (D) Bar graph comparing the apparent viscosity [Pa·s] at each cell stage, calculated from the MSD data in (B); Error bars represent 95% confidence intervals.

### Establishing magnetic tweezer-based active microrheology using ferrofluid probes in early *C. elegans* embryos

While PMR revealed a decrease in probe mobility during embryogenesis (Fig. 1B,C), it does not distinguish whether this reflects changes in intracellular driving forces or in the mechanical properties of the cytoplasm, such as increased viscosity. To directly assess cytoplasmic mechanics, we implemented active microrheology (AMR) using magnetic tweezers. To enable AMR measurements in living embryos, we used a magnetic fluid (ferrofluid) as a deformable probe (He et al., 2014; Doubrovinski et al., 2017; Orii and Tanimoto, 2025). We succeeded to injected ferrofluid into the *C. elegans* gonad, where it formed spherical droplets of a few micrometers in diameter (Fig. 2A, top). After 3.5–6.5 h of incubation, these droplets were incorporated into embryos through normal oogenesis (Fig. 2A, bottom). This establishes a method for generating micrometer-sized probes within living *C. elegans* embryos.

**Figure 2.**
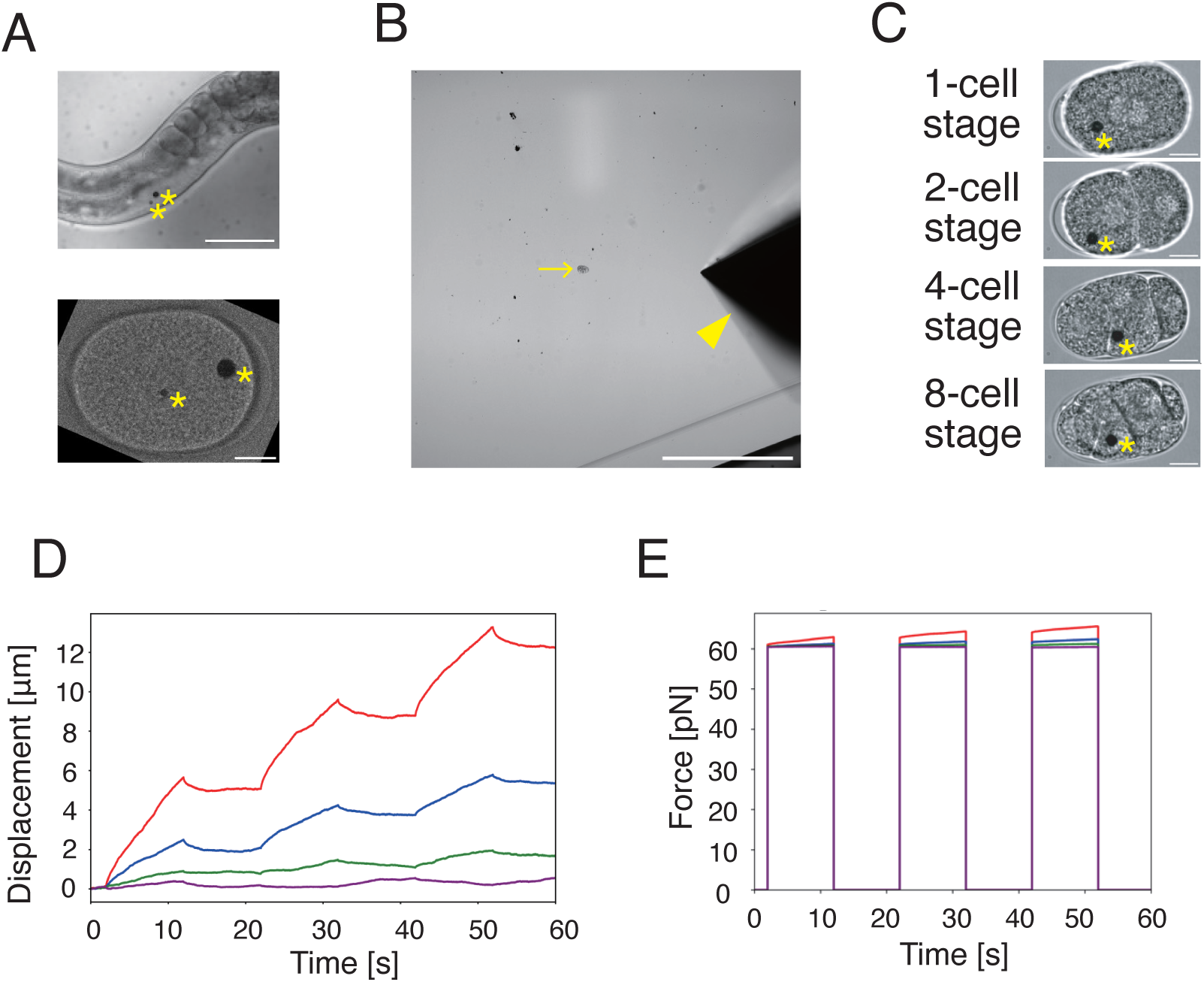
Using ferrofluid droplets to perform active microrheology (AMR) in *C. elegans* embryos. (A) (Top) Bright-field image of an adult *C. elegans* hermaphrodite with magnetic fluid introduced into the gonad. Scale bar: 100 μm. (Bottom) Bright-field image of a 1-cell stage *C. elegans* embryo containing a ferrofluid droplet, observed approximately 4.5 hours after the gonadal introduction shown in (Top). The location of the ferrofluid is marked by a yellow asterisk. Scale bar: 10 μm. (B) Bright-field image illustrating the spatial arrangement of the *C. elegans* embryo and the magnetic tweezer. The embryo is indicated by a yellow arrow, and the magnetic tweezer is indicated by a yellow arrowhead. Scale bar: 200 μm. (C) Bright-field images of *C. elegans* embryos from the 1-cell to the 8-cell stage containing ferrofluid droplets used for AMR with magnetic tweezers. Scale bar: 10 μm. (D–E) Relative changes in the x-coordinate position of droplets (D) and applied force (E) over time in the embryos shown in (C). Data are color-coded by cell stage: 1-cell stage (red), 2-cell stage (blue), 4-cell stage (green), and 8-cell stage (purple).

For AMR measurements, we selected droplets of approximately 4 μm in diameter (3.62 ± 0.34 μm, mean ± SD), as larger droplets (5–10 μm) were frequently retained in the gonad and rarely incorporated into embryos. Droplets were subjected to repeated force-application cycles consisting of a 10 s loading phase followed by a 10 s relaxation phase. During loading, droplets were translated along the embryonic long axis at a target speed of ∼0.1 μm/s, comparable to the timescale of intracellular movements such as nuclear migration (Kimura and Onami., 2005) (mean speed = 0.52 μm/s, 1 cell stage), from positions sufficiently distant from the cell boundary, ensuring that they did not reach the cortex during displacement.

From the displacement trajectories of the droplets (Fig. 2D) and force calibrations (Fig. 2E, Supplementary Fig. S1), We quantified the mechanical response of the cytoplasm by calculating the creep compliance *J*(*t*) (Fig. 3A). This quantity was obtained by normalizing the measured displacement by the applied magnetic force and probe size, based on the generalized Stokes relation (see Methods), thereby providing a measure of the intrinsic mechanical properties of the cytoplasm. The creep compliance *J*(*t*) describes how easily a material deforms over time in response to a constant force. In this framework, higher compliance indicates that the material deforms more readily (i.e., is mechanically softer), whereas lower compliance indicates greater resistance to deformation.

**Figure 3.**
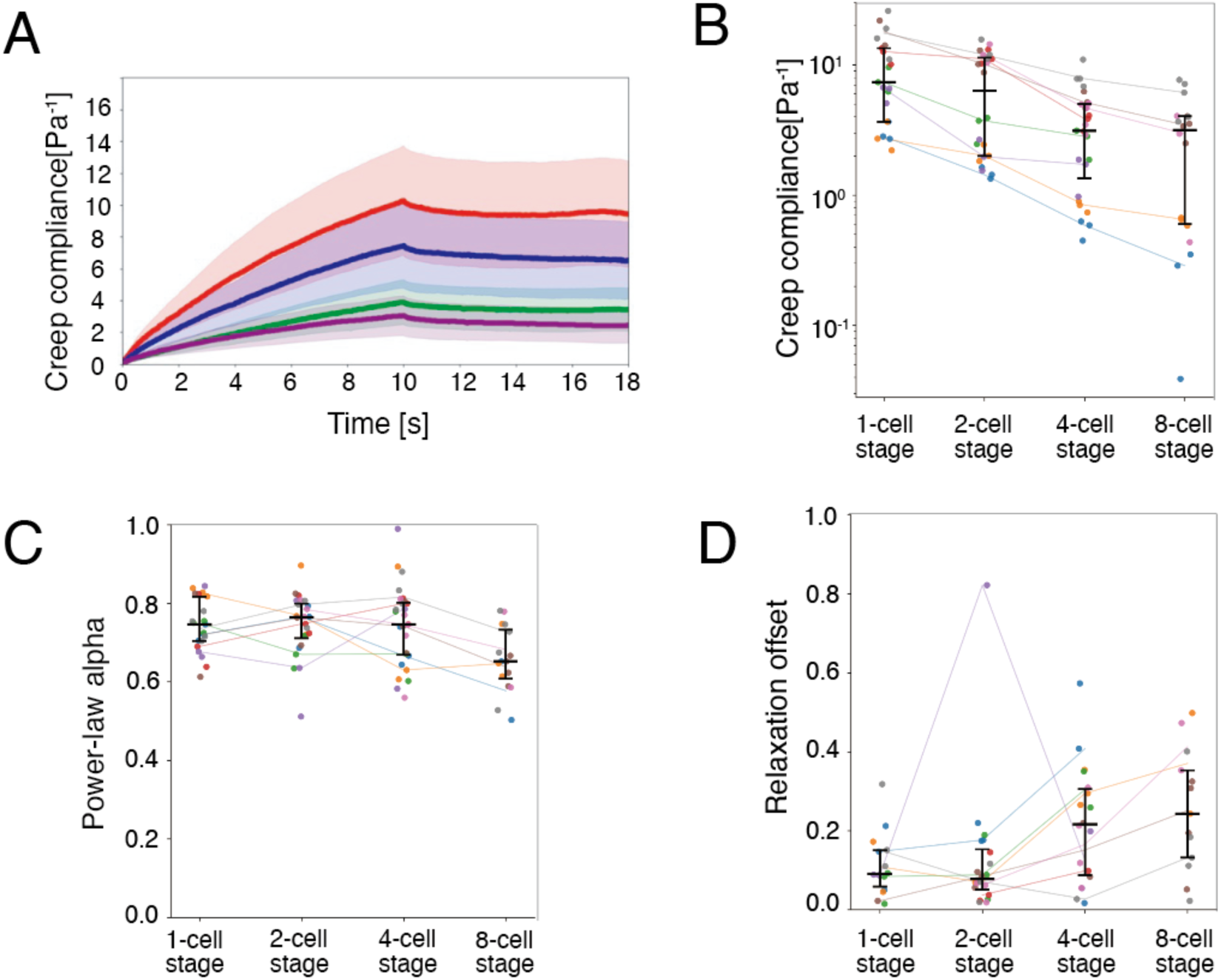
Active microrheology (AMR) characterized the viscoelasticity of cytoplasm in *C. elegans* embryos. (A) Time-dependent creep compliance measured at each cell stage on the original time scale. Color coding for embryo stages: 1-cell stage in red, 2-cell stage in blue, 4-cell stage in green, 8-cell stage in purple. (B) Creep compliance values at 10 s after force application plotted for each cell stage. (C) Power-law exponents obtained by fitting the time-dependent creep compliance during the loading phase for each cell stage. (D) Recovery ratio during the relaxation phase, calculated as the percentage of deformation recovered after force release, plotted for each cell stage. In (B)–(D), individual embryos are distinguished by color, and values are connected across cell stages using the median for each embryo at each stage. Black lines indicate the group median and interquartile range (IQR) at each cell stage. Sample sizes were: 1-cell stage, N = 7 embryos and n = 21 trajectories; 2-cell stage, N = 8 and n = 24; 4-cell stage, N = 8 and n = 27; 8-cell stage, N = 5 and n = 18.

The measured compliance at 1 s in the 1-cell stage (2.1 ± 1.5 Pa^-1^) corresponds to an apparent viscosity of ∼0.5 Pa·s, under the Newtonian-fluid assumption (Supplementary Table S2). The estimated viscosity is approximately three orders of magnitude higher than that of water. This estimate is also close to the viscosity independently obtained by PMR using 40 nm beads (∼0.30 Pa·s in the 1-cell stage; Fig. 1C), supporting the consistency between passive and active microrheology measurements in the same system. The estimated viscosity is also comparable to that reported for the cytoplasm of early *Drosophila* embryos measured by ferrofluid-based AMR (∼1.5 Pa·s) (Doubrovinski et al., 2017), despite the difference in species. In addition, the measured value is similar to the viscosity reported for *C. elegans* P granules (∼1 Pa·s), which was estimated independently from droplet deformation under shear stress (Brangwynne et al., 2009). Together, these comparisons indicate that our measurements capture cytoplasmic mechanical properties within the expected range.

### Active microrheology reveals a progressive decrease in cytoplasmic compliance during early embryogenesis

To enable quantitative comparison across developmental stages, we focused on matched cell-cycle stages: prometaphase at the 1-cell stage and interphase at later stages (see Methods). We found that creep compliance *J*(*t*) decreased progressively as embryogenesis proceeded from the 1-cell to the 8-cell stage (Fig. 3A–B). For droplets of the same size, compliance was highest at the 1-cell stage and declined steadily through the 2-, 4-, and 8-cell stages (Supplementary Table S3). At 10 s, the mean compliance in the 8-cell stage was 3.5-fold lower than in the 1-cell stage, corresponding to a 72% reduction. These results indicate that the cytoplasm becomes progressively more resistant to deformation during early embryogenesis, consistent with the results obtained from PMR using 40-nm beads (Fig. 1B-D).

The creep compliance exhibited power-law scaling over the 10-s observation window, with an exponent of 0.7 across developmental stages (Fig. 3C), indicating that viscous behavior remains dominant within this timescale. Upon removal of the applied force, deformation recovered only partially (Fig. 3E), further supporting a viscoelastic response with predominant viscous dissipation and a measurable elastic component. The apparent elastic modulus estimated from the recovered compliance showed no clear stage-dependent change (Fig. S2B; Supplementary Table S2). Taken together, these results indicate that the developmental decrease in compliance is mainly attributable to increased cytoplasmic viscosity.

### Linear mixed-effects modeling confirms stage-dependent stiffening during early embryogenesis

Creep compliance decreased progressively during embryogenesis (Fig. 3). However, substantial variability in creep compliance was observed even within each developmental stage (Fig. 3). To determine whether this variability could be explained by measurement conditions and whether the stage-dependent trend remained after accounting for such factors, we analyzed the data using a linear mixed-effects model.

We considered probe size and intracellular position as potential sources of variability. Previous studies in sea urchin embryos (Najafi et al., 2023) and *Dictyostelium* (Wilhelm, 2008) have shown that larger probes tend to yield higher apparent viscosity and elasticity, and that when probes are comparable in size to the cell, the measured viscoelastic properties can depend on intracellular position, particularly on the distance from the cell membrane (Najafi et al., 2023). In addition, because multiple measurements were obtained from the same embryo, embryo-to-embryo differences were also expected to contribute to the observed variability.

We therefore fitted a linear mixed-effects model (LMM) to creep compliance at 10 s, including developmental stage, probe size, and intracellular position as fixed effects (Fig. 4A; Supplementary Table S4), and embryo identity as a random intercept. Intracellular position was decomposed into components parallel and perpendicular to the direction of droplet motion. Specifically, for observation *i* from embryo *j* at cell stage *k*, the creep compliance at 10 s, *y_ijk_*, was modeled as

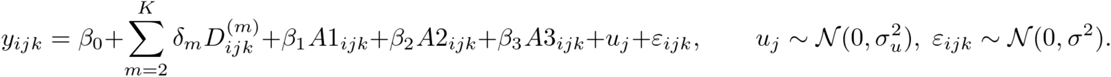

**Figure 4.**
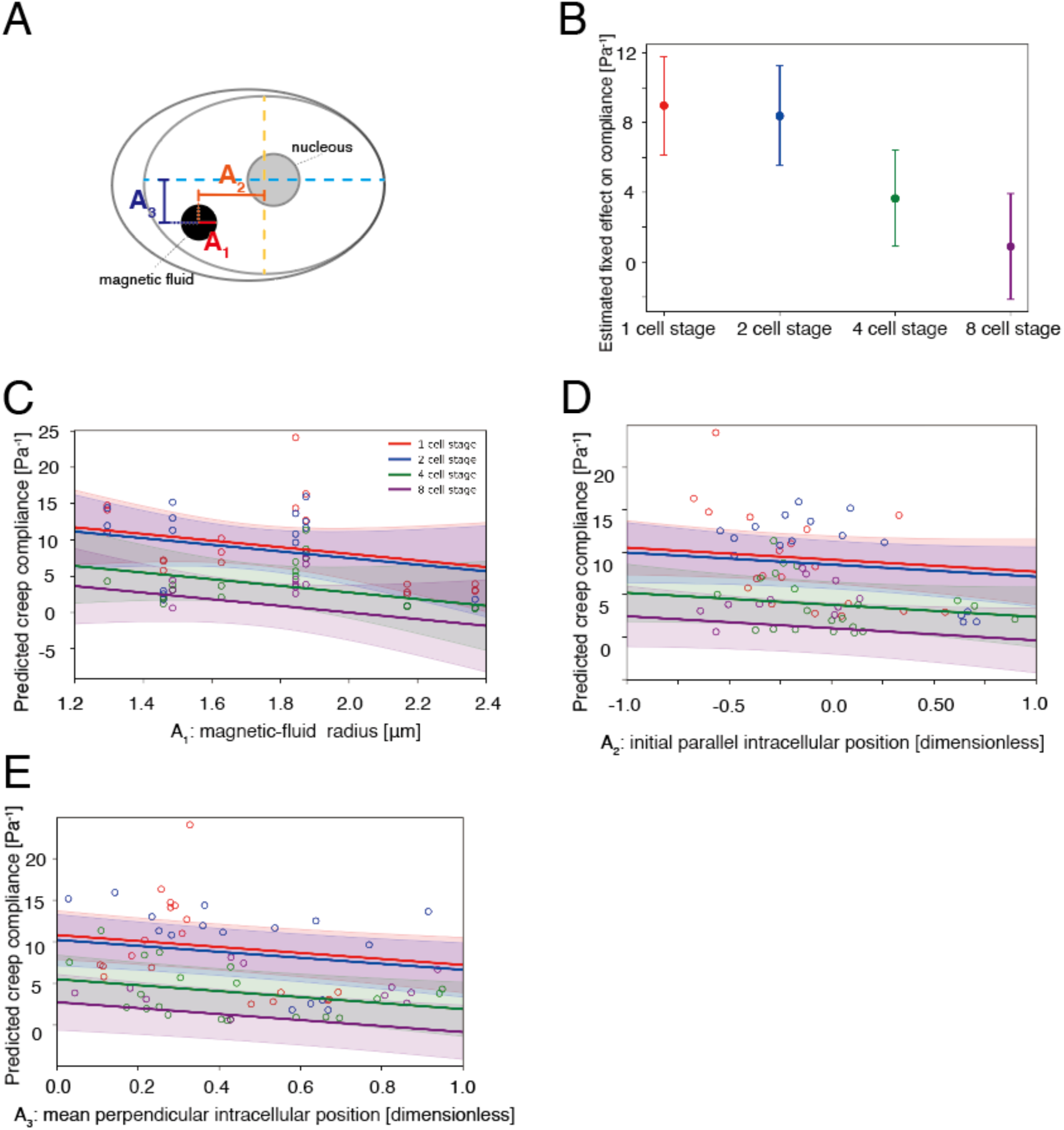
Fixed and partial effects of developmental stage and measurement-related variables on creep compliance estimated by the linear mixed-effects model. (A) Schematic representation of the three continuous covariates considered in the linear mixed-effects model: magnetic droplet radius (A1), initial relative position parallel to the direction of droplet motion (A2), and mean absolute relative position perpendicular to the direction of motion (A3). (B) Fixed-effect estimates for each developmental stage are shown as stage-specific values from the linear mixed-effects model: the intercept for the 1-cell stage (reference level) and the intercept plus the corresponding stage coefficient for the other stages. Points indicate estimated values and error bars indicate 95% confidence intervals. (C-E) Predicted creep compliance is plotted as a function of (C) magnetic droplet radius (A1), (D) initial intracellular position parallel to the direction of droplet motion (A2), and (E) mean intracellular position perpendicular to the direction of droplet motion (A3), shown on their original measurement scales. Colored lines indicate model predictions for each developmental stage (1-cell, red; 2-cell, blue; 4-cell, green; 8-cell, purple), and shaded regions indicate the corresponding 95% confidence intervals. Open circles show observed data points from the corresponding stage.

Here, *A1*_ijk_ is magnetic droplet radius; *A2*_ijk_ is the initial relative position parallel to the direction of droplet motion; *A3*_ijk_ is the mean relative position perpendicular to the direction of motion; and *D*^(m)^_ijk_ is a stage dummy variable, with one stage treated as the reference level. *β*_0_ is the intercept, *β*_m_ represents the fixed effect of stage m relative to the reference stage, and *β*_1_, *β*_2_, and *β*_3_ are regression coefficients for the continuous covariates. *u_j_* is the embryo-level random intercept, assumed to follow a normal distribution with variance *σ*_u_^2^, and *ε_ijk_* is the residual error term, assumed to follow a normal distribution with variance *σ*^2^.

Even after accounting for these factors, creep compliance remained significantly lower at the 4-cell and 8-cell stages compared to the 1-cell stage, with a similar but non-significant trend at the 2-cell stage (Fig. 4B; Supplementary Table S5). Thus, the developmental decrease in cytoplasmic compliance is robust to variation in measurement conditions.

Among the measurement-related variables, probe size showed a negative but non-significant association with creep compliance (Fig. 4C; see Discussion). Intracellular position parallel to the direction of motion had no statistically significant effect (Fig. 4D). In contrast, the perpendicular component showed a significant negative association with compliance, indicating that droplets located closer to the cell periphery exhibited lower creep compliance (Fig. 4E). Finally, substantial embryo-to-embryo variability remained after accounting for these factors. The estimated random-intercept variance exceeded the residual variance (Supplementary Table S6; intraclass correlation coefficient, ICC = 0.73). This indicates that 73% of the variability arose from differences between embryos. Despite this variability, the stage-dependent decrease in compliance was consistently detected, indicating that the observed mechanical changes reflect a robust feature of cytoplasmic organization during early embryogenesis.

### Cytoplasmic stiffening occurs independently of cytokinesis

Having established that cytoplasmic compliance decreases during early embryogenesis, we next asked whether this change reflects intrinsic developmental regulation or geometric effects associated with decreasing cell size. Previous studies have shown that probe mobility depends on cell size and proximity to the cell boundary (Najafi et al., 2023), both of which change during cleavage, the observed decrease could in principle be explained by cell size reduction.

To distinguish between these possibilities, we disrupted cytokinesis by RNAi-mediated knockdown of the *zen-4* gene, which encodes a kinesin essential for cleavage furrow formation (Powers et al., 1998; Davies et al., 2014). Under these conditions, the cell cycle progressed and nuclear division occurred, but cytokinesis failed, resulting in a single, multinucleated cell that did not physically divide (Fig. 5A). If the decrease in cytoplasmic compliance is driven by cell size, it should be abolished under these conditions; if not, the decrease should persist even in the absence of cytokinesis.

**Figure 5:**
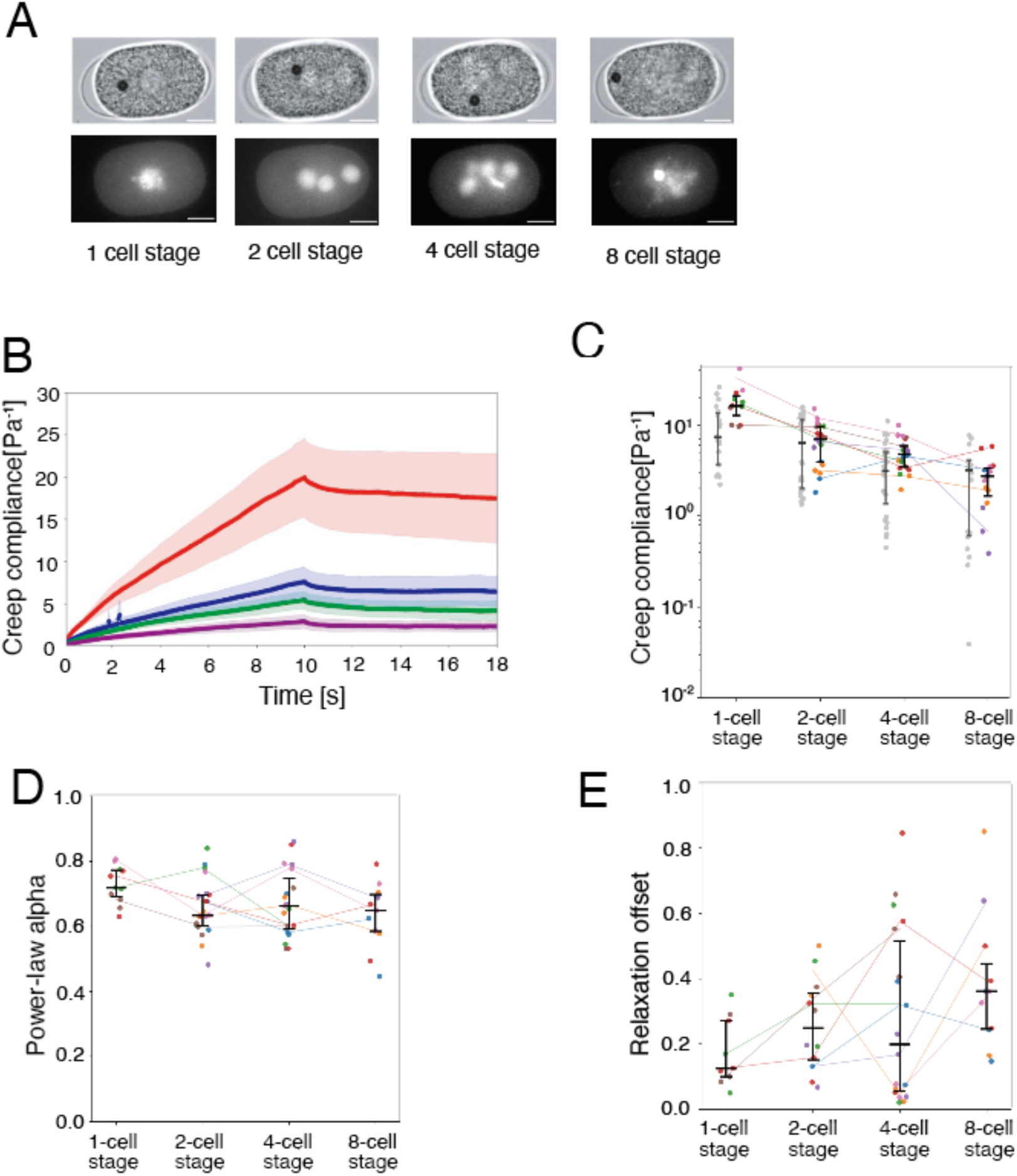
Time-dependent creep compliance in *zen-4* RNAi embryos across developmental stages. (A) Representative bright-field images (top) and fluorescence images (bottom) of zen-4 RNAi embryos at the 1-, 2-, 4-, and 8-cell stages. (B) Time-dependent creep compliance measured at each cell stage on the original time scale. Color coding for embryo stages: 1-cell stage in red, 2-cell stage in blue, 4-cell stage in green, 8-cell stage in purple. (C) Creep compliance values at 10 s after force application plotted for each cell stage. Mutant embryos are distinguished by color, while WT embryos are shown in gray on the left side of each stage; the WT data are the same as those presented in Fig. 3C. (D) Power-law exponents obtained by fitting the time-dependent creep compliance for each cell stage. Individual embryos are distinguished by color. (E) Recovery ratio during the relaxation phase, calculated as the percentage of deformation recovered after force release, plotted for each cell stage. In (C)–(E), individual embryos are distinguished by color, and values are connected across cell stages using the median for each embryo at each stage. Black lines indicate the group median and interquartile range (IQR) at each cell stage.Sample sizes were: 1-cell stage, N = 4 embryos and n = 11 trajectories; 2-cell stage, N = 7 and n = 21; 4-cell stage, N = 7 and n = 21; 8-cell stage, N = 5 and n = 15.

Strikingly, despite the absence of cytokinesis, the creep compliance of micrometer-sized probes decreased over time (Fig. 5B, C, Supplementary Table S3). Although the compliance values at the 1- and 4-cell stages were slightly higher under *zen-4* RNAi, the magnitude and temporal profile of the decrease were comparable to those observed in wild-type embryos. These results demonstrate that the developmental decrease in cytoplasmic compliance does not require physical cell division and cannot be explained by simple mechanical consequences such as reduced cell size or increased probe–membrane interactions. Instead, it reflects an intrinsic, actively regulated program of cytoplasmic remodeling during early embryogenesis.

## Discussion

We found that intracellular mobility decreases progressively during early embryogenesis in *C. elegans*. Both passive and active microrheology (PMR and AMR) revealed consistent reductions in particle motion across length scales, indicating that local cytoplasmic environment becomes progressively less compliant during development.

A key technical advance of this study is the implementation of ferrofluid-based AMR in *C. elegans* embryos, enabling quantitative measurements of cytoplasmic mechanical properties throughout early embryogenesis. Previous studies in this system employed PMR(Daniels et al., 2006) or AMR of specific intracellular structures such as the mitotic spindle using 1- or 2.8- *μ* m magnetic particles (Garzon-Coral et al., 2016). In contrast, the ferrofluid-based approach used here allows the formation and manipulation of micrometer-sized droplets directly within the cytoplasm, enabling more general characterization of the mechanical environment experienced by large intracellular objects. Because droplet size can be varied over a micrometer range, this method also expands the accessible measurement scales compared with previous bead-based approaches. Furthermore, by applying defined external forces, AMR enables direct assessment of cytoplasmic mechanical properties independently of endogenous intracellular driving forces (Hoffman et al., 2006; Mizuno et al., 2007; Wilhelm, 2008). Combined with PMR measurements using 40-nm beads, this approach allowed us to characterize developmental changes in cytoplasmic mechanics across distinct observational scales during embryogenesis.

Although both PMR and AMR consistently revealed a developmental decrease in probe mobility during embryogenesis, several differences were also observed between the two measurements. AMR measurements using ∼4 μm droplets exhibited a smaller power-law exponent (∼0.7, Fig. 3C) than PMR measurements using 40-nm beads (∼1, Fig. 1D), indicating a relatively large elastic contribution for the larger probe. In addition, the apparent viscosity estimated from AMR (Supplementary Table S2) was several-fold higher than that estimated from PMR (Fig. 1D) at comparable developmental stages. These differences may reflect the distinct physical principles underlying the two methods, as PMR infers mechanical properties from spontaneous fluctuations under assumptions related to thermal equilibrium, whereas AMR directly measures force-induced mechanical responses. Alternatively, they may arise from the large difference in probe size between the two approaches. Consistent with the latter possibility, linear mixed-effects analysis within the AMR dataset suggested a negative association between probe size and creep compliance (Fig. 4C), although the effect did not reach statistical significance. Future studies combining PMR and AMR using comparable probe sizes will help disentangle the respective contributions of probe size and measurement modality to intracellular mechanical measurements.

A major conceptual implication of this study is that local cytoplasmic mechanical properties are dynamically regulated during development. The observed decrease in compliance over successive cleavage divisions indicates a progressive reduction in local cytoplasmic compliance at the probe scale over a short developmental timescale. These findings suggest that local intracellular mechanics are not merely passive constraints, but are actively modulated as part of the developmental program.

This change may provide a unifying physical framework for diverse embryonic phenomena. During embryogenesis, the size and dynamics of many intracellular features change gradually, including reductions in the sizes of nuclei and mitotic spindles (Hara and Kimura, 2009; Wühr et al., 2008; Good et al., 2013; Crowder et al., 2015; Lacroix et al., 2018), centrosome size (Greenan et al., 2010; Decker et al., 2011), as well as decreases in the rates of spindle elongation (Hara and Kimura, 2009; Okafornta et al., 2025) and nuclear growth (Fickentscher et al., 2024), increased resistance to displacement of the mitotic spindle (Garzon-Coral et al., 2016), and elongation of cell-cycle duration (Arata et al., 2015; Fickentscher et al., 2018), in *C. elegans* and other organisms. These observations have largely been interpreted in terms of geometric constraints associated with decreasing cell size, for example through the reduced availability of cytoplasmic components required to build intracellular structures. Our results suggest that changes in the material state of the cytoplasm may also contribute to these phenomena. The observed decrease in compliance persists even when cell size is uncoupled from division, arguing against a purely geometric explanation. A progressively less compliant local mechanical environment could limit the growth and movement of large intracellular structures such as nuclei and spindles, potentially contributing to their reduced size and slower dynamics. In addition, cytoplasmic viscosity can influence biochemical activity (Fukuhara et al., 2025). More broadly, changes in cytoplasmic material properties have been shown to modulate diffusion-controlled processes and intracellular reaction dynamics in living cells, including microtubule polymerization and depolymerization (Persson et al., 2020; Molines et al., 2022). Therefore, changes in cytoplasmic mechanics may affect intracellular transport and reaction rates, with possible consequences for broader cellular organization. Thus, the progressive decrease in cytoplasmic compliance may contribute to the coordinated changes in cellular state that accompany early embryogenesis, providing a shared physical context that complements geometric explanations.

The molecular and physical mechanisms underlying this regulation remain to be determined. Possible contributors include cytoskeletal reorganization and active remodeling, changes in molecular crowding, cytoplasmic phase separation and condensate formation, and nonequilibrium processes driven by molecular motors. Recent studies have highlighted that the cytoplasm behaves as an active, heterogeneous material whose mechanical state influences organelle positioning, intracellular transport, and reaction dynamics (Eskandari et al., 2025; Arjona et al., 2023). Consistent with this view, cytoplasmic mechanics are likely shaped by the interplay of multiple factors, including cytoskeletal networks (Fabry et al., 2001; Hurst et al., 2021; Muenker et al., 2024; Orii and Tanimoto, 2025; Kickuth et al., 2026; Nishi et al., 2026), membrane organization (Hiramoto, 1970; Merta et al., 2021), colloidal packing (Joyner et al., 2016; Delarue et al., 2018), and ATP-dependent active forces (Guo et al., 2014; Ebata et al., 2023). Understanding how these factors are coordinated during development will be an important direction for future research. Clarifying the mechanisms that regulate cytoplasmic mechanics may provide new insights into how cellular organization and dynamics are controlled through the material state of the cytoplasm.

## MATERIALS AND METHODS

### C. elegans strains

*C. elegans* strains were maintained at 22°C on nematode growth medium (NGM) plates seeded with OP50 *Escherichia coli*, as described previously (Brenner, 1974). Strain names and genotypes are provided in Supplemental Table S1.

### Preparation of PLL-g-PEG‒coated fluorescent beads

To reduce nonspecific surface interactions, 40-nm carboxylate-modified fluorescent beads (FluoSpheres, red-orange, Thermo Fisher Scientific, F8794) were coated with PLL-g-PEG following Torisawa et al. (2025). Beads were mixed with PLL-g-PEG solution (100 μg/mL in 1× PBS, SuSoSu), briefly sonicated using an ultrasonic cleaner (Branson 1510), and incubated with rotation for 1 h at room temperature.

After coating, beads were washed with 1× PBS (Takara T900) using centrifugal filters (Amicon Ultra, 0.5 mL, 100 kDa cutoff, Merck, UFC510024) to remove unbound PLL-g-PEG. The final bead suspension was adjusted to the desired concentration in 1× PBS and stored at 4 °C until use.

### Microinjection of PLL-g-PEG‒coated FluoSpheres and ferrofluid into *C. elegans* gonads

Young adult hermaphrodites (N2 strain, maintained at 22 °C) were mounted on 2% agarose pads on glass coverslips (24 × 55 mm, Matsunami) and covered with halocarbon oil 700 (Sigma). Gonads were visualized using an inverted microscope (Zeiss Axiovert), and injection solutions were introduced using pulled glass microneedles.

For passive microrheology (PMR), PLL-g-PEG‒coated FluoSpheres were injected using a FemtoJet microinjector (Eppendorf) and microneedles prepared with a P-1000 pipette puller (Sutter Instrument). Prior to injection, bead suspensions were centrifuged to remove aggregates, and approximately 2 μL of the supernatant was loaded into the injection needle.

For active microrheology (AMR), ferrofluid (DS-60, Sigma HiChemical) was injected using a manual microinjection system consisting of a CellTram 4r Oil injector and an InjectMan 4 micromanipulator (Eppendorf), as described previously (Orii and Tanimoto, 2025). Injection pipettes were prepared from thin-walled borosilicate glass capillaries (TW100-4, WPI) using a P-1000 puller. For FluoSphere injection, the FemtoJet was operated at an injection pressure of p_i = 1000 hPa, an injection time of t_i = 0.2 s.

After injection, worms were released from the agarose pads by adding 5 μL of M9 buffer and transferred to 35 mm NGM plates seeded with OP50 bacteria. Animals injected with FluoSpheres were incubated at 22 °C for at least 3 h before imaging, whereas those injected with ferrofluid were incubated for at least 3.5 h.

### Imaging of *C. elegans* embryos

Embryos were prepared for imaging as described previously (Torisawa and Kimura, 2022). Briefly, adult worms were dissected in 0.75× egg salt buffer (118 mM NaCl, 40 mM KCl, 3.4 mM CaCl₂, 3.4 mM MgCl₂, 5 mM HEPES, pH 7.2). Isolated embryos were placed on 2% agarose pads on glass slides (76 × 26 mm, Matsunami) with spacers, covered with M9 buffer, and sealed with an 18 × 18 mm coverslip (Matsunami).

Time-lapse imaging was performed using a spinning-disk confocal microscope consisting of an inverted microscope (Olympus IX71) equipped with a Yokogawa CSU-MP unit. Images were acquired using a 60× silicone-immersion objective (Olympus, UPlanSApo, 60×/1.30Sil) with an additional 2.0× intermediate magnification. An EM-CCD camera (Andor iXon) controlled by NIS Elements software (Nikon) was used for acquisition.

Embryos were first inspected in bright-field to confirm normal morphology. Fluorescence imaging was performed using 561-nm excitation at full laser power. Unless otherwise specified, time-lapse images were acquired for 2,000 frames at 30 ms intervals. For the CAL2895 strain, continuous imaging was performed from the 1-cell stage for 60,000 frames under the same conditions. Under these conditions, the effective pixel size was 0.136 μm/pixel, the exposure time was 30 ms per frame, and the excitation laser was operated at 100% power.

### Particle tracking and trajectory analysis

Particle trajectories were extracted using Fiji and analyzed with custom software (Mark2, Furuta & Toyoshima, 2008). A 4-frame moving average was applied to smooth trajectories. To minimize contamination by aggregated particles, tracer spots were selected by visual inspection, excluding obvious clusters or non–point-like signals. For each embryo at each developmental stage, approximately 50 trajectories were analyzed; with N=5 embryos per stage, this yielded ∼250 trajectories per stage (at least >200 trajectories).

### Magnetic tweezers setup

AMR measurements were performed using home-built electromagnetic tweezers with feedback control (Orii and Tanimoto, 2024, Miyamoto et al., 2026). Briefly, the core of magnetic tweezers was cylindrical magnetic materials of 8 mm diameter made from 45-Permalloy (Nilaco,782624). One end of the core was tapered to achieve a tip radius of ∼2 um. The magnetic core was inserted into a solenoid coil with 1000 windings of copper wires (AWG 24 or 26). The electromagnetic tweezers were held by a micro-manipulator mounted on a motorized microscope (Ti2-E, Nikon) equipped with a CFI Plan Apo 40x Lambda DIC N2 objective (MRD00405, Nikon), 1.5x intermediate magnification, a LED light source (X-Cite XYLIS, excelitas), and a Digital CMOS camera (ORCA-Fusion, HAMAMATSU). The magnetic tweezers and camera were synchronized using a trigger signal from HCImage software (Hamamatsu Photonics).

### Calibration of magnetic tweezers using ferrofluid droplets

Magnetic force calibration was performed using ferrofluid droplets, following established approaches (Tanimoto et al., 2018; Orii and Tanimoto, 2025). Briefly, ferrofluid droplets within the size range used in experiments were dispersed in a glycerol solution of known viscosity (η = 0.22 Pa·s at 20 ℃).

Droplet motion under an applied magnetic field was recorded, and steady-state velocities were measured. The magnetic force acting on each droplet was calculated using Stokes’ law, F = 6π η R v, where η is the viscosity of the medium, R is the droplet radius, and v is the measured velocity.

The force‒distance relationship of the magnetic tweezers was determined by measuring forces at different distances (x) from the tweezer tip. The resulting data were fitted to an empirical function of the form:

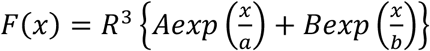

where R is the droplet radius, *x* is the distance from the tweezer tip, F is the magnetic force, a and b are decay lengths, and A and B are fitting coefficients.

Fitting yielded the following parameter values: A = 78.5 (pN/μm^3^), a = 168 (μm), B = 3.54 (pN/μm^3^), and b = 7.19 (μm). These fitted parameters were subsequently used to estimate the magnetic force applied to droplets in embryos based on their distance from the tweezer tip.

### Creep experiments

For creep experiments, two magnetic tweezers were initially positioned at opposite ends along the longitudinal axis of the embryo. Ferrofluid droplets were first translated toward one end of the embryo by applying a magnetic field. The position of the magnetic tweezers tip was fixed at a distance of 600 ∼ 800 μm from the sample during a measurement. Magnetic field was approximately 7.2 mT at the blunt end of the tweezers.

After positioning, one of the magnetic tweezers was removed, and creep measurements were performed using the remaining tweezer. A constant magnetic field was applied for 10 s (loading phase), followed by a 10 s period without magnetic field (relaxation phase); this sequence was defined as one cycle. Each measurement consisted of one to three consecutive cycles. Each measurement consisted of three consecutive cycles. At the onset of loading, the droplet–tweezer distance was 499 µm (range, 437–561µm), corresponding to an estimated force of 28.6 pN (range, 6.0–65.6 pN).

These measurements were repeated in the same embryo from the stage prior to nuclear fusion through the 8-cell stage.

### Creep compliance analysis

Creep compliance J(t) was calculated based on the generalized Stokes relation, assuming that the ferrofluid droplet behaves as a spherical probe embedded in a linear viscoelastic medium and that inertial effects are negligible under low-Reynolds-number conditions (Furst and Squires, 2017).

For the loading phase (response phase), creep compliance was calculated as

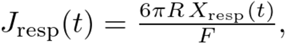

Where

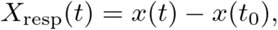

*x(t)* is the droplet position, t_0_ is the onset of force application, F is the applied magnetic force, and R is the droplet radius.

For the unloading phase (relaxation phase), the analysis was performed under the additional assumption that the principle of linear superposition holds. Under this assumption, force release was treated as a negative step in the applied load, and relaxation compliance was calculated as

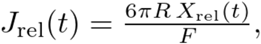

where

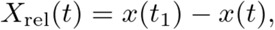

and t_1_ is the onset of unloading. In this definition, X_rel_(t) represents the recovery displacement measured from the droplet position at the moment of force release. To estimate the apparent viscosity, we used the creep compliance measured 1 s after the onset of force application. Under the Newtonian-fluid assumption, creep compliance is proportional to time as:

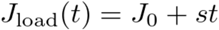

where J_0_ is the intercept and s is the slope of the creep compliance. The slope s corresponds to the rate of increase in compliance, and the apparent viscosity was estimated as the inverse of this slope:

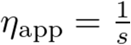

This value represents an apparent viscosity estimated from the short-time creep response within the first 1 s of force application.

To estimate the apparent elastic component, we quantified the amount of compliance recovered during the relaxation phase (Ziemann et al., 1994). The compliance at 10 s during the loading phase was defined as J_load,end_, and the compliance remaining at the end of the relaxation phase was defined as J_relax,end_. Specifically, J_elax,end_ was calculated as the median compliance value over the final 20% of the relaxation phase.

The recovered elastic compliance was then calculated as

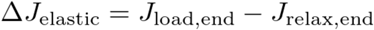

and the apparent elastic modulus was estimated as the inverse of this recovered compliance:

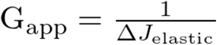

This value represents an apparent elastic modulus based on the recoverable deformation within the experimental observation window.

### Data fitting and parameter estimation

Mean squared displacement (MSD) data (Figure 1B–D) were analyzed as follows. MSD curves were first fitted to a power-law model, MSD = A *t^α^*, to assess the scaling exponent α. Because α was close to 1 over the fitting range (Fig. 1D), indicating approximately normal diffusion, we then estimated the diffusion coefficient D by fitting the same MSD data to the linear relation MSD = 4Dt (2D). Apparent viscosity was calculated from D using the Stokes–Einstein equation, η = k_B_ T / (6π a D), where η is viscosity, k_B_ is the Boltzmann constant, T is absolute temperature, a is the bead radius, and D is the diffusion coefficient.

AMR data were analyzed by calculating the creep compliance J(t) as described above and fitting it to a power-law function, J(t)=B t^β^, where B is the prefactor and β is the scaling exponent.

All curve fitting was performed in Python using the SciPy curve_fit function for nonlinear least-squares optimization. Fits were performed over lag times of 0.03–1 s for PMR and 0.005–10 s for AMR. No weighting was applied unless otherwise specified.

### Linear mixed-effects modeling

Creep compliance at 10 s, J(10 s), was analyzed using a linear mixed-effects model. For observation i from embryo j at cell stage k, the response variable *y_ijk_* was modeled with cell stage as a fixed effect and embryo identity as a random intercept.

The following continuous covariates were included as fixed effects: (1) magnetic droplet radius (μm), (2) initial relative position parallel to the direction of droplet motion (dimensionless, range [-1, 1]), and (3) mean absolute relative position perpendicular to the direction of motion (dimensionless, range [0, 1]).

All continuous covariates were z-standardized prior to analysis to improve numerical stability and enable comparison of effect sizes across variables. Standardized coefficients therefore represent the expected change in J(10 s) associated with a one standard deviation increase in each covariate.

Cell stage was encoded using dummy variables, with one stage treated as the reference level. Embryo-to-embryo variability was modeled using a random intercept, and both random effects and residual errors were assumed to follow normal distributions.

To facilitate interpretation on the original measurement scale, standardized coefficients were additionally converted to per-unit effects by dividing by the corresponding standard deviation of each covariate. Standard errors and confidence intervals were transformed accordingly.

Model adequacy was assessed using standard diagnostic procedures. Observed values were compared with model-predicted values to evaluate overall fit (Fig. S3A). Conditional residuals (including both fixed and random effects) were examined by plotting standardized residuals against fitted values to detect systematic deviations (Fig. S3B). Normality of residuals was assessed using quantile‒quantile plots (Fig. S3C).

**Table 1:** Fixed-effect estimates from the linear mixed-effects model. The model was fitted to creep compliance measured 10 s after force application. Developmental stage was included as a categorical fixed effect with the 1-cell stage as the reference level. Magnetic droplet radius (*A1*), intracellular position parallel to droplet motion (*A2*), and intracellular position perpendicular to droplet motion (*A3*) were included as continuous fixed effects, and embryo identity was modeled as a random intercept. For continuous predictors, coefficients are shown both per 1 SD increase and per 1 unit increase.

| Term | Coef. per 1<br>SD | SE per 1<br>SD | z | p | 95% CI<br>lower | 95% CI<br>upper |
| --- | --- | --- | --- | --- | --- | --- |
| Intercept | 8.97 | 1.44 | 6.24 | <0.001 | 6.15 | 11.78 |
| 2 cell stage | -0.59 | 0.90 | -0.65 | 0.51 | -2.35 | 1.17 |
| 4 cell stage | -5.32 | 0.76 | -6.98 | <0.001 | -6.82 | -3.83 |
| 8 cell stage | -8.08 | 1.07 | -7.58 | <0.001 | -10.17 | -5.99 |
| A1 | -1.46 | 1.34 | -1.09 | 0.28 | -4.07 | 1.16 |
| A2 | -0.52 | 0.40 | -1.30 | 0.19 | -1.30 | 0.26 |
| A3 | -0.91 | 0.37 | -2.48 | 0.01 | -1.63 | -0.19 |

**Table 1 (continued)**
| Term | SD of x | Coef. per 1 unit | SE per 1 unit | 95% CI<br>lower | 95% CI<br>upper |
| --- | --- | --- | --- | --- | --- |
| A1 | 0.34 | -4.25 | 3.89 | -11.88 | 3.38 |
| A2 | 0.39 | -1.34 | 1.03 | -3.36 | 0.68 |
| A3 | 0.26 | -3.47 | 1.40 | -6.21 | -0.73 |

## Acknowledgments

We thank Ms. Tomoko Ozawa and Ms. Nozomi Takase for their assistance with the experiments, the members of the Cell Architecture Laboratory, Physics and Cell Biology Laboratory of National Institute of Genetics, and the Tanimoto Lab of Yokohama City University for discussion and advice. Some strains were provided by the CGC, which is funded by NIH Office of Research Infrastructure Programs (P40 OD010440). This work was supported by JST SPRING Grant Number JPMJSP2104 for SK, and by JSPS KAKENHI Grant Numbers 25K02276 for AK, 24K09405 for TT.

---

**Supplementary Figure S1.**
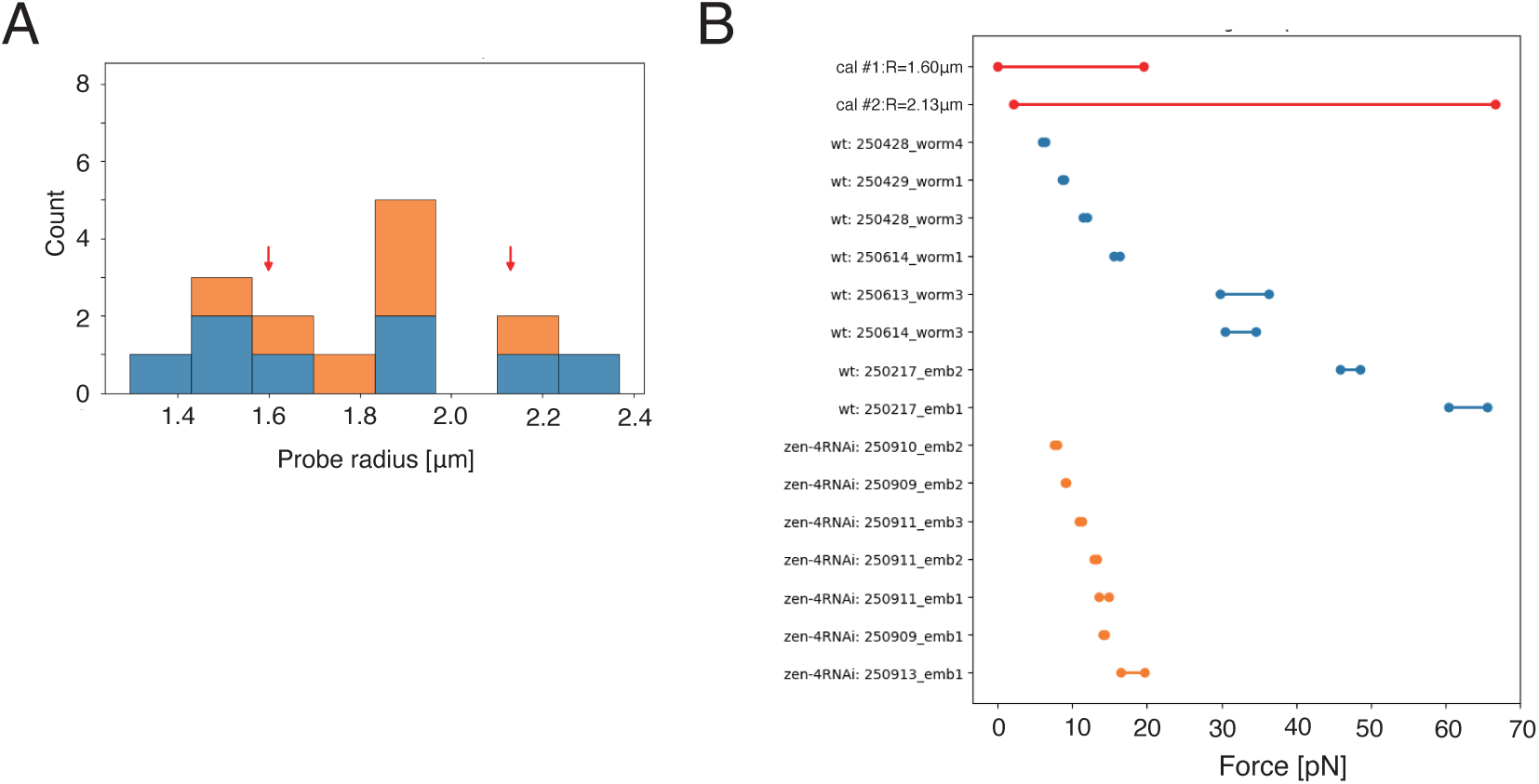
Comparison of probe-size distribution and force range between calibration and experimental datasets. (A) Histogram of probe radii used in the experimental datasets. Probe sizes from the control (wt, blue) and *zen-4* RNAi (orange) datasets are shown as stacked histograms, where each individual was counted once because probe radius was constant within an individual. The two probe radii used for calibration are indicated by two red arrows. (B) Comparison of force ranges between the calibration and experimental datasets. For the two calibration datasets, the ranges between the minimum and maximum forces are shown with red lines. For each individual in the experimental datasets, the force range during the applied-force intervals is shown with blue (control) and orange (*zen-4* RNAi) lines. These plots were used to assess whether the current calibration datasets reasonably cover the probe-size range and force range of the experimental data. Since the probe sizes were comparable and the force ranges largely overlapped, the current calibration was considered to be broadly applicable to the present analysis without substantial extrapolation.

**Supplementary Figure S2.**
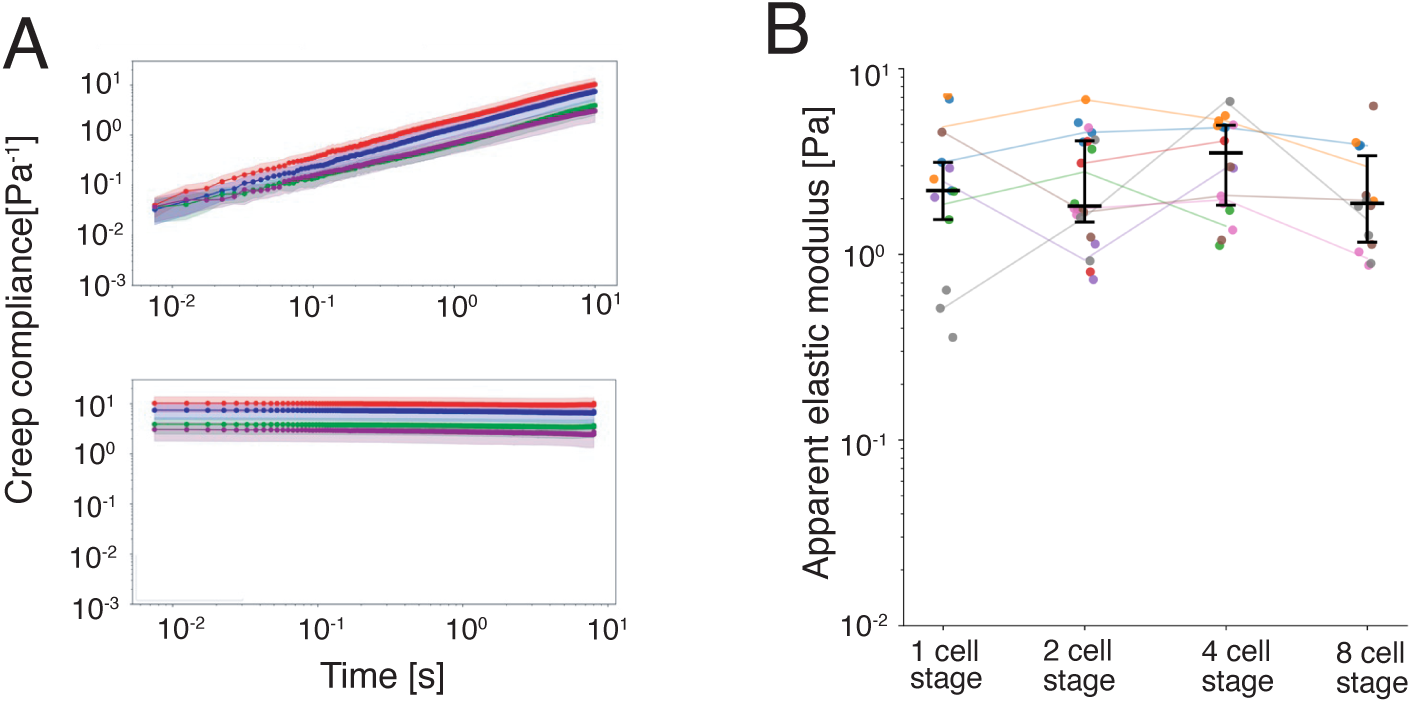
Time-dependent creep compliance and recovery-derived apparent elasticity. (A) Time-dependent creep compliance shown in Fig. 3A plotted on a logarithmic time scale. The loading phase is shown on the top, and the relaxation phase is shown on the bottom. Embryo stages are color coded as follows: 1-cell stage, red; 2-cell stage, blue; 4-cell stage, green; 8-cell stage, purple. (B) Apparent elastic modulus estimated from the compliance recovered during the relaxation phase. For each segment, the recovered elastic compliance was defined as the difference between the compliance at the end of loading and the compliance at the end of relaxation, and the apparent elastic modulus was calculated as its inverse. Data are plotted for each cell stage, individual embryos are distinguished by color, and values are connected across cell stages using the median for each embryo at each stage. Black lines indicate the group median and interquartile range (IQR) at each cell stage.

**Supplementary Figure S3.**
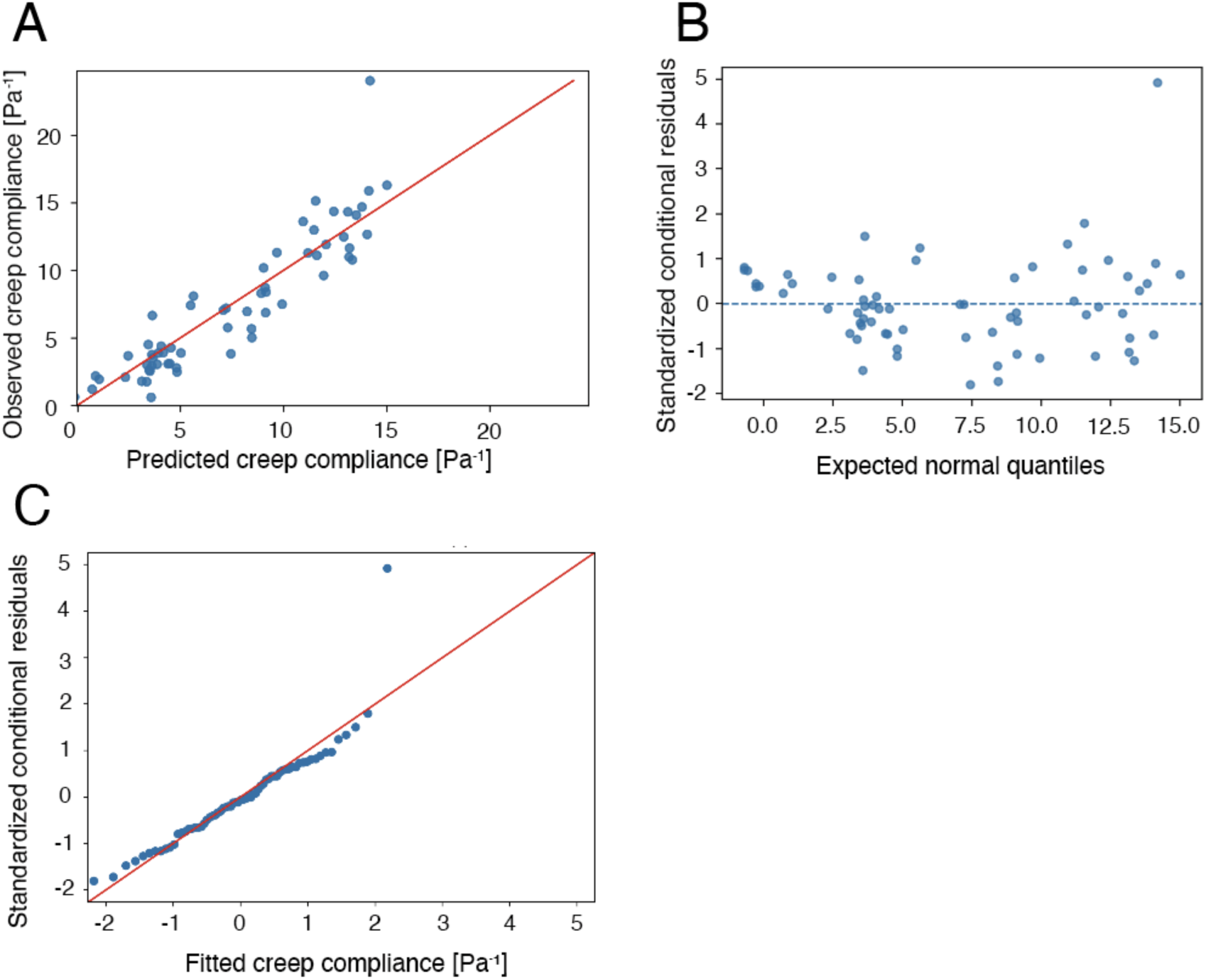
Diagnostic assessment of the linear mixed-effects model. (A) Observed versus model-predicted creep compliance. Blue points indicate individual observations, and the red solid line indicates the identity line. (B) Standardized conditional residuals plotted against fitted values. Conditional residuals were defined as the differences between the observed creep compliance values and the fitted values from the full mixed-effects model, including both fixed effects and embryo-specific random intercepts. (C) Quantile–quantile (Q–Q) plot of the conditional residuals.

## Notes

### Competing Interest Statement

The authors have declared no competing interest.

